# Topography of Hippocampal Connectivity with Sensorimotor Cortex Revealed by Optimizing Smoothing Kernel and Voxel Size

**DOI:** 10.1101/2020.05.14.096339

**Authors:** Douglas D. Burman

## Abstract

Studies of the hippocampus use smaller voxel sizes and smoothing kernels than cortical activation studies, typically using a multivoxel seed with specified radius for connectivity analysis. This study identified optimal processing parameters for evaluating hippocampal connectivity with sensorimotor cortex (SMC), comparing effectiveness by varying parameters during both activation and connectivity analysis. Using both 3mm and 4mm isovoxels, smoothing kernels of 6-10mm were evaluated on the amplitude and extent of motor activation and hippocampal connectivity with SMC. Psychophysiological interactions identified hippocampal connectivity with SMC during volitional movements, and connectivity effects from multivoxel seeds were compared with alternate methods; a structural seed represented the mean connectivity map from all voxels within a region, whereas a functional seed represented the regional voxel with maximal SMC connectivity. With few exceptions, the same parameters were optimal for activation and connectivity. Larger isovoxels showed larger activation volumes in both SMC and the hippocampus; connectivity volumes from structural seeds were also larger, except from the posterior hippocampus. Regardless of voxel size, the 10mm smoothing kernel generated larger activation and connectivity volumes from structural seeds, as well as larger beta estimates at connectivity maxima; structural seeds also produced larger connectivity volumes than multivoxel seeds. Functional seeds showed lesser effects from voxel size and smoothing kernels. Optimal parameters revealed topography in structural seed connectivity along both the longitudinal axis and mediolateral axis of the hippocampus. These results indicate larger voxels and smoothing kernels improve sensitivity for detecting both cortical activation and hippocampal connectivity.

## Introduction

Noise can obscure the weak blood oxygen-level dependent (BOLD) response used to detect activation and functional connectivity in functional magnetic resonance imaging (fMRI). A number of approaches have been tried over the years to improve signal detection. Two fundamental considerations are voxel size and the size of the smoothing kernel (1). Optimal voxel size should generally be commensurate with activation volume, although due to susceptibility field gradients, smaller voxel sizes may sometimes increase sensitivity despite less total signal intensity (2, 3). Larger voxel sizes improve signal strength but provide lower spatial resolution, whereas hippocampal studies generally prefer better resolution due its small size; during analysis, voxel dimensions within any plane typically range from 1mm to 3mm (4–12).

Spatial smoothing is used to increase signal-to-noise in BOLD signals and provide smoothness to imaged data (13, 14). Smoothing expands the size of detected activation by reducing noise; generally, the optimal smoothing kernel is 2-3 times the voxel size, both for individual analysis (15, 16) and for large group analysis, with larger smoothing kernels suggested for group analysis in studies with fewer subjects (17). Smoothing kernels may degrade the resolution of signal (18), so some studies of the hippocampus avoid smoothing (12), whereas others use a small Gaussian smoothing kernel ranging from 3-6mm full width at half maximum (FWHM) (6, 8, 10, 19, 20). Nonetheless, several hippocampal studies have used larger smoothing kernels of 8mm (4, 5, 9, 11, 21).

The selection of appropriate processing parameters raises challenges for connectivity studies, especially those designed to examine the influence of hippocampal activity on cortical areas that differ in optimal processing parameters. Generally, the optimal smoothing kernel for connectivity analysis is 2-3 times the voxel size (22), as it is for activation analysis. With few exceptions (23, 24), smoothing kernels used in hippocampal connectivity studies are small, ranging from 2-6mm (25).

The effects of voxel and smoothing kernel size in the hippocampus have not been explored empirically, so their precise relationship to activation and connectivity is unknown. Although effective in activating the hippocampus and its connections with prefrontal cortex (PFC), memory tasks may be less than optimal to identify effects of processing parameters, as the actual size, intensity, and location of memory-related PFC activation may depend on memory content (26–31). The sensorimotor cortex (SMC), by contrast, shows robust activation based on movements of the represented body part, known from studies of topography (32–35); furthermore, a recent study showed hippocampal-SMC connectivity restricted to the hand representation during a hand movement task (36).

Building on that work, the current study systematically explores the effect of voxel size, smoothing kernel, and seed selection on task-specific activation and connectivity from hippocampus to sensorimotor cortex (SMC). A larger voxel size (4mm vs. 3mm isovoxels) generated a modest increase in cortical activation and hippocampal connectivity volumes, except from posterior hippocampus, whereas larger smoothing kernels consistently increased both cortical activation and connectivity, particularly connectivity from structural seeds. Structural seeds, representing the mean connectivity from all voxels in a specified region of the hippocampus, proved superior to conventional multivoxel seeds, providing larger connectivity volumes and a demonstrable topographic organization.

## Materials and methods

### Data availability

Subjects, task, data acquisition, and processing procedures have previously been described in detail (36, 37), with the dataset available for download (38).

### Subjects

Thirteen normal right-handed adults participated in the study (ages 24-59, mean=42.3, five females) following written consent; informed consent procedures complied with the Code of Ethics set forth in the Declaration of Helsinki, and were approved by the Institutional Review Board at the NorthShore University HealthSystem / Research Institute.

### Experimental task

Each subject performed a visual/motor task, consisting of 6 cycles of a specified sequence of visual and motor conditions over a period of 6m 4s. Each cycle included a block of sequential tapping and a block of repetitive tapping, each separated by a passive visual condition. The task ended with 10s of passive fixation on a central cross.

During the sequential tapping block, a 4-button sequence to be remembered was displayed while a metronome ticked at 2 Hz; the subject pressed the remembered sequence of buttons in time with the metronome once the onscreen instructions were replaced with a cross. Subjects repeated the 4-button sequence throughout the 16s block, which ended when a circular checkerboard pattern flickered onscreen for 9s. During this visual block, subjects fixated the center of the pattern and refrained from moving.

During the repetitive tapping block, the subject was instructed to tap the same finger on both hands in synchrony with the metronome. Once the instruction screen was replaced with the number ‘1’, the subject tapped the index finger from both hands; every 4s, the onscreen number increased by one, and the subject changed finger. This motor condition was also followed by the passive visual condition.

Button presses were recorded from the right hand during both motor tasks to verify accurate performance. Behavioral analysis demonstrated motor learning and recall effects during the sequence learning task; anticipatory movements prior to the metronome demonstrated cognitive control of movements during both repetitive tapping and sequence learning (36).

### MRI data acquisition

Images were acquired using a 12-channel head coil in a 3 Tesla Siemens scanner. Blood-oxygen level dependent (BOLD) functional images were acquired with the echo planar imaging (EPI), using the following parameters: time of echo (TE) = 25 ms, flip angle = 90°, matrix size = 64 × 64, field of view = 22×22 cm, slice thickness = 3 mm, number of slices = 32; time of repetition (TR) = 2000 ms; and the number of repetitions = 182. These parameters produced in-plane voxel dimensions of 3.4 × 3.4mm. A structural T1 weighted 3D image (TR = 1600 ms, TE = 3.46 ms, flip angle = 9°, matrix size = 256 × 256, field of view = 22×22 cm, slice thickness = 1 mm, number of slices = 144) was acquired in the same orientation as the functional images.

### fMRI data processing

Data were processed and analyzed using SPM12 software (http://www.fil.ion.ucl.ac.uk/spm). Images were spatially aligned to the first volume to correct for small movements. After removing one cycle from a subject’s data set due to excessive head movements, maximal RMS movement for any subject was less than 1.5mm (mean = 0.850 ± 0.291mm). Sinc interpolation minimized timing-errors between slices; functional images were coregistered to the anatomical image, normalized to the standard T1 Montreal Neurological Institute (MNI) template, and resliced for two sets of analyses. For one dataset, functional images were resliced as 3mm isovoxels, potentially improving in-plane image resolution by incorporating differences in signal intensity associated with small head movements. For the other dataset, functional images were resliced as 4mm isovoxels, potentially increasing sensitivity while decreasing image resolution.

To identify task-related activation, copies of images from each dataset were smoothed with a 6mm, 8mm, or 10mm isotropic Gaussian kernel and filtered with a high pass cutoff frequency of 128s. Conditions of interest were specified for sequence learning, visual, and repetitive tapping, then modeled for block analysis using a canonical hemodynamic response function.

### Activation analysis

A parameter estimate of the BOLD response to each condition was generated; motor activation was identified for each subject by contrasting mean BOLD responses to motor vs. passive visual conditions. Analyses used an intensity threshold of p<0.05 with a family-wise error (FWE) correction for multiple comparisons, applied to a sensorimotor region of interest (ROI). Typically, this ROI was specified as the overlap between TD and aal atlas labels for post- plus precentral gyrus in the WFUPickatlas toolbox for SPM. An additional analysis used the hand representation as the ROI (36) in order to evaluate whether effects of kernel size and smoothing kernel had a similar effect on activation parameters within the region known to be active.

For each voxel size and smoothing kernel combination, the mean and standard error of activation parameters from all individual subjects were recorded and quantitatively compared; these parameters included t-value threshold (corresponding to that required by the FWE correction), maxima t-values, and contrast amplitudes. Additionally, paired t-tests applied to individual subject data allowed statistical evaluation of different voxel sizes with each smoothing kernel, and of different smoothing kernels with the same voxel size.

For analysis of group effects, BOLD contrasts from individual subjects during the motor memory and repetitive tapping conditions were entered into a 1-sample t-test for each voxel size / smoothing kernel combination, recording the same parameters as for individuals (t-threshold, maxima t-values, and contrast amplitudes). In addition, paired t-tests looked for significant group differences in activation between 6mm and 10mm smoothing kernels.

Activation maps from different conditions were overlapped to allow visual comparisons for differences in the size and location of activation.

### Connectivity analysis

#### PPI analysis

Psychophysiological interactions (PPI) were used to identify task-specific connectivity of the hippocampus with sensorimotor cortex (39–41). The general approach was described previously (36). For 4mm isovoxels, 78 voxels were identified from the left hippocampus of the normalized brain and 78 from the right, as delimited by the aal atlas in the WFU PickAtlas toolbox (http://fmri.wfubmc.edu/software/PickAtlas); each seed was wholly contained within the hippocampus. For 3mm isovoxels, 278 voxels were identified in the left hippocampus.

At each hippocampal voxel, a contrast was selected and specified to create eigenvariates for all conditions in the statistical model; an interaction term then specified greater effect on activity from the motor condition than from the visual condition. After adjustments for regional differences in timing and baseline activity, a regression analysis showed the magnitude of the BOLD signal that correlated with this interaction term elsewhere in the brain. The magnitude of SMC connectivity was quantified from each hippocampal voxel.

#### Seed selection

Single- and multivoxel approaches were both used to select seeds for connectivity analysis. A 3×3 matrix was created by dividing the hippocampus into thirds of equal distance along the A-P plane, then dividing each section into equal thirds across the medial-lateral plane. Because the head of the hippocampus is enlarged, anterior seeds contained more voxels than middle or posterior seeds, the number of voxels in each structural seed was the same across the medial-lateral plane. Anatomical seeds were labelled by their position within the matrix (A to C from anterior to posterior, 1 to 3 from medial to lateral) and the sign of connectivity (positive or negative).

A “structural” seed’s connectivity map was calculated as the mean connectivity from all voxels within the region. Alternatively, a single “multivoxel” seed was selected from the center of the anatomical location with a 6mm radius; this provided comparable spatial evaluation of connectivity from 3mm isovoxels (a diameter of 4 voxels) and 4mm isovoxels (a diameter of 3 voxels). To minimize asymmetry effects, comparisons in connectivity across voxel sizes were limited to the left hippocampus.

Single-voxel seeds could be based either on anatomical location or function. Anatomical selection was based on spaced intervals within the structural seeds, providing finer-grain analysis that was useful for examining topography. “Functional” seed selection was restricted to voxels within (or adjacent to) a structural seed showing significant connectivity; the single voxel was selected that showed maximal connectivity within the SMC mask. By reflecting individual variability in functional localization, functional seeds sought to provide a better estimate of the magnitude and extent of hippocampal influence on SMC activity. Functional seeds were identified from the left and right hippocampus for both sequence learning (“memseed”) and repetitive tapping (“tapseed”), using global analysis from both sides to improve sensitivity (36).

#### PPI Group analysis

For the sequence learning task, beta estimates of connectivity from each subject’s left hippocampus were entered into a 1-sample random effects analysis; the hand representation delineated from activation analysis was used as the ROI, applying an intensity threshold of p<0.05 with FWE correction. A separate analysis was run for multivoxel and structural seeds at 4 combinations of voxel size (3mm and 4mm) with smoothing kernel (6mm and 10mm).

To improve sensitivity for repetitive tapping, beta estimates of connectivity from each subject’s left and right hippocampus were entered into an ANOVA model for global analysis; as before, the hand representation delineated from activation analysis was used as the ROI. This ROI provides optimal sensitivity for detecting effects, as hippocampal connectivity with SMC during volitional finger movements is selective for the hand representation (36).

## Results

### Effects of voxel size and smoothing kernels on SMC activation

Group and individual analyses showed similar trends of voxel size and smoothing kernels on patterns of SMC activation, as described below. During group analyses, differences in group activation between 3mm and 4mm isovoxels appeared minor, as were most smoothing kernel effects *except* for activation volume. At the individual level, however, effects from different voxel sizes and smoothing kernels were both significant

For both 3mm and 4mm isovoxels, maxima values generally decreased incrementally with larger smoothing kernels. The maximal t-value decreased (Table 1, Fig 1A), a pattern observed during both group and individual analyses; lower maxima t-values were observed during group analysis. Estimates for the magnitude of contrast effects also decreased, especially for individual analyses; the magnitude of effects nearly converged for individual and group analyses for 10mm smoothing (see bottom of graphs). In addition, the threshold to identify significant activation dropped, resulting from less local variability in signal amplitude with increasing kernel size. During both individual and group analyses, the net effect was larger activation volumes with larger smoothing kernels, with 4mm isovoxels always generating larger activation volumes than 3mm isovoxels (Fig 1B). With a 10mm smoothing kernel, group differences in SMC activation between the two voxel sizes overlapped extensively, but with a slightly larger activation volume for 4mm isovoxels during sequence learning (Fig 1C).

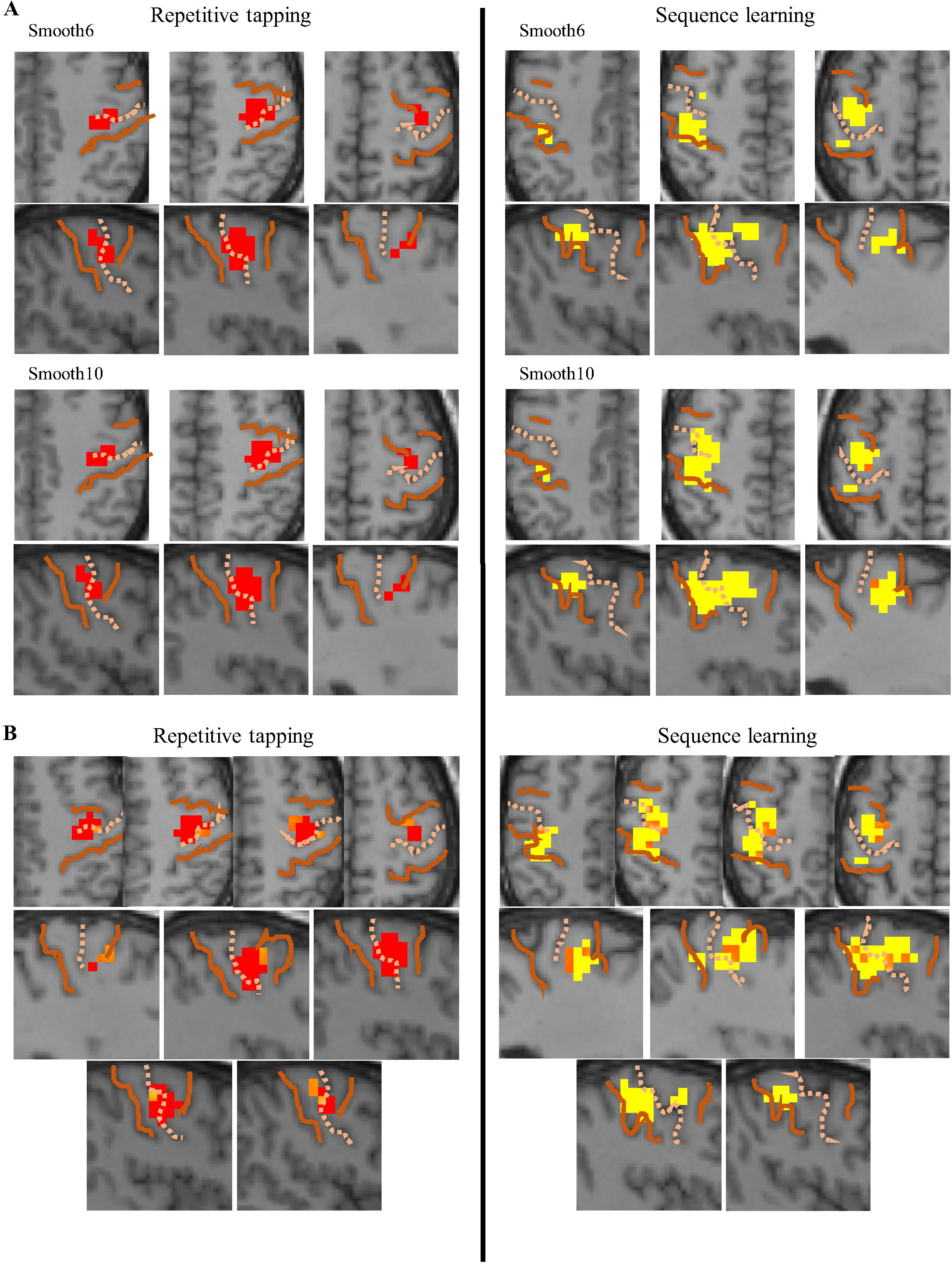

Larger smoothing kernels thus provide greater sensitivity for detecting activation by reducing noise (random variability in signal). Comparing changes in activation volume with different ROIs provides further insight (Table 1). With the SMC mask, large increases in activation volume were observed as the smoothing kernel increased 6mm to 10mm, volume increases of 84.2 - 440.0%. These increases were much smaller within the hand representation, never exceeding 33.3% and nearly non-existent for 3mm isovoxels (volume increases of 1.9 – 1.6% for repetitive tapping and sequence learning, respectively). This suggests larger smoothing kernels are particularly effective for improving statistical detection by reducing signal variability outside the activated region, thus requiring a smaller mean difference in signal to detect activation.

Because data from the same individuals were processed with different voxel size and smoothing kernels, differences in activation parameters resulting from different processing methods could be measured directly for each individual. The standard error for mean differences in parameters across the group of subjects often differed greatly for both repetitive tapping and sequence learning; nonetheless, paired t-tests showed that individuals consistently had lower activation thresholds and greater activation volumes with larger (4mm) isovoxels, regardless of smoothing kernel (Table 2).

For each task, the effects of different smoothing kernels on activation parameters were also evaluated with paired t-tests during each motor task (Tables 3-4). With 3mm isovoxels, larger smoothing kernels during both motor tasks generated higher activation volumes, but significantly lower maxima t-values, contrast magnitudes, and activation thresholds. Often, these patterns were also observed with 4mm isovoxels; however, no differences were observed between 8mm and 10mm smoothing kernels for activation threshold during repetitive tapping (Table 3), or for the maxima t-values and contrast magnitudes during sequence learning (Table 4).

To summarize, 4mm isovoxels provided greater sensitivity to activation than 3mm isovoxels (lower threshold and higher activation volume), as did 10mm compared with 6mm smoothing kernels (for both 3mm and 4mm isovoxels). For 4mm isovoxels, differences between 8mm and 10mm smoothing kernels were less differentiated for some activation parameters.

### Effect of smoothing kernels on hippocampal motor activation

Using the hippocampal ROI mask, group analysis failed to demonstrate motor activation for any combination of voxel size and smoothing kernel. Using paired t-tests to compare activation following 10mm vs. 6mm smoothing, however, more intense activation was observed from the larger smoothing kernel during both motor tasks (Fig 2). For both 3mm and 4mm isovoxels, the most intense activation (orange) was observed along the border between structural seeds B1 and C1; mildly elevated activation (yellow) was observed throughout B1.

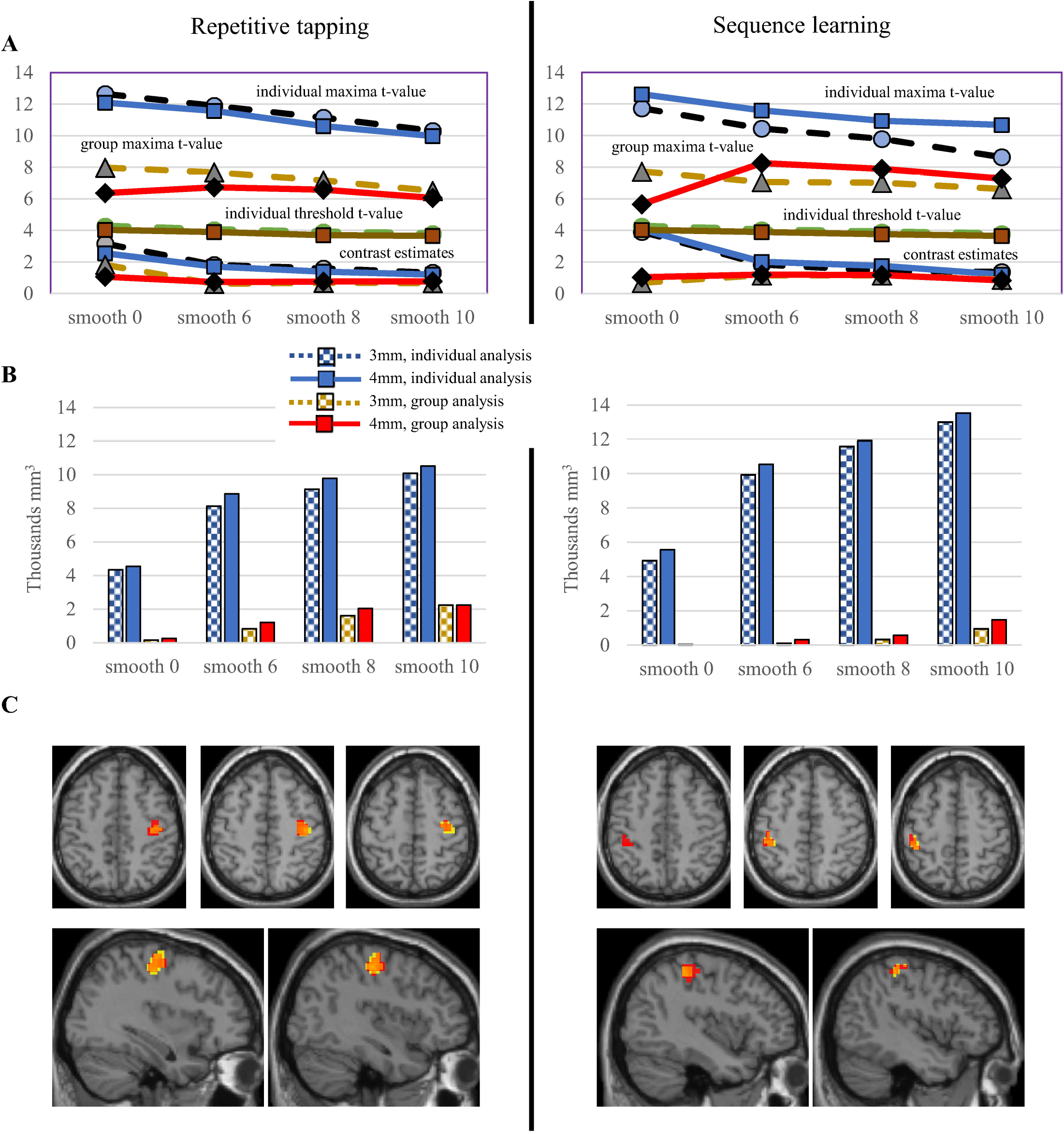

### Hippocampal connectivity: Structural seeds

#### Effects of voxel size and smoothing on hippocampal connectivity

Inverse connectivity from many regions of the hippocampus was observed during sequence learning (Fig 3 and Table 5). For a given combination of voxel size and smoothing kernel, a structural seed (striped and dotted bars) generated a larger volume of SMC connectivity than the corresponding multivoxel seed (dashed and solid bars), even when the parameter estimate at the postcentral (black dot) or precentral maximum (red dot) was larger for the multivoxel seed. In some cases, differences in methodology could determine whether connectivity remained significant following an additional correction for testing these nine regions (stars). The connectivity from A1 and A2 seeds, for example, were significant only for structural seeds derived from 4mm isovoxels with 10mm smoothing.

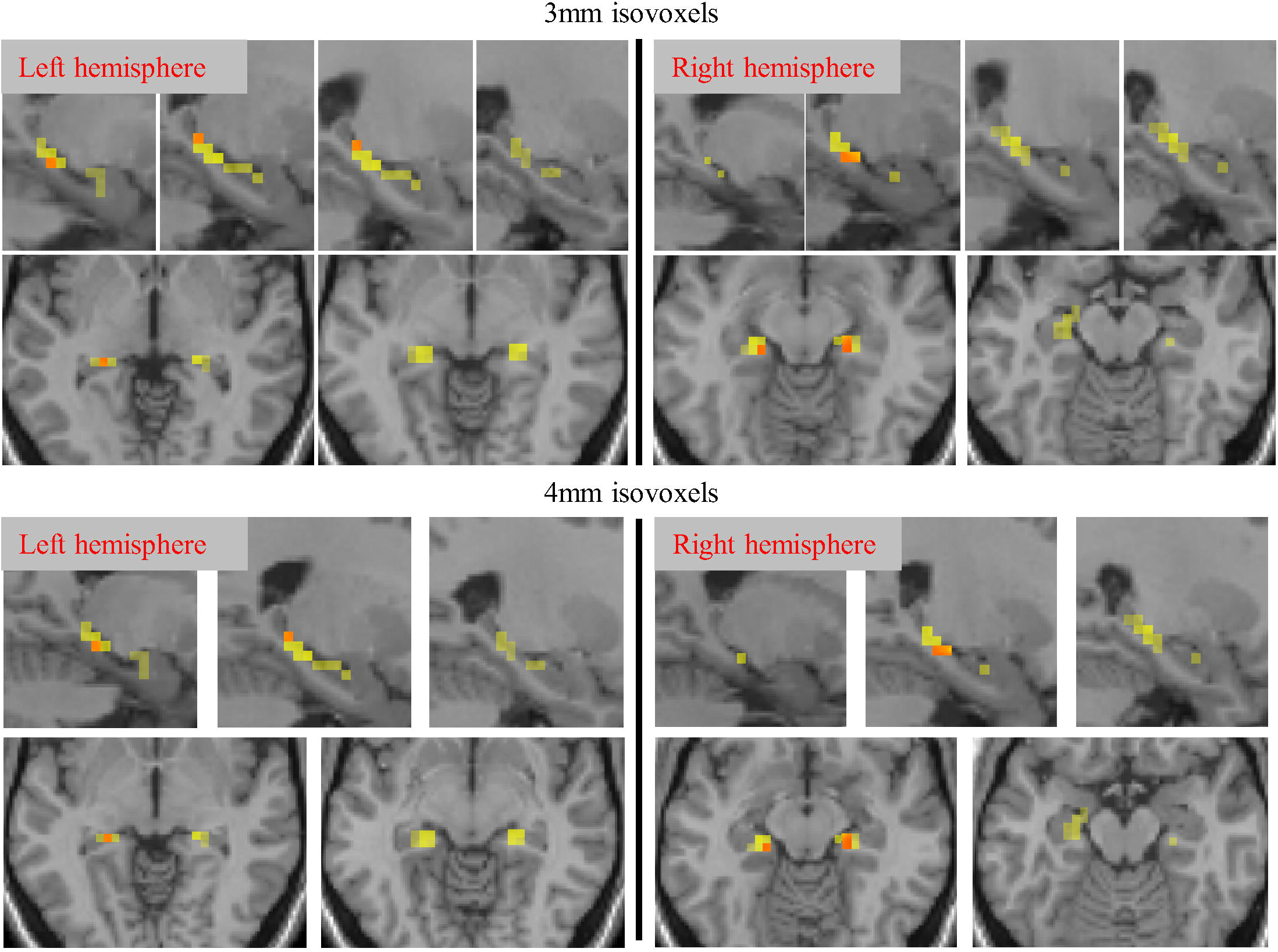

In anterior (A1-A3) and middle hippocampus (B1-B2), the volume of connectivity during random effects analysis was typically greater for 4mm isovoxels, both for structural and multivoxel seeds; only in posterior hippocampus (C1-C3) was the volume of connectivity greater for 3mm isovoxels (see also Table 6). In paired t-tests, beta parameter estimates in structural seeds were significantly larger for 4mm voxels and also with the larger smoothing kernel (10mm), especially at postcentral maxima (Table 6). Overall, the volume of connectivity was larger for structural seeds than multivoxel seeds (invariably so when effects survived the FWE correction), even when maxima parameter estimates were significantly smaller (observed only with the smaller 6mm smoothing kernel).

Connectivity demonstrated through different methodologies overlapped extensively, particularly when the volume of connectivity was similar (see Fig 3, seed B1- at bottom left). When one method generated a much larger volume, however, an additional locus of connectivity could appear (in bottom right of Fig 3, see precentral connectivity for S3 method at seed C1-). The volume of connectivity thus reflected the sensitivity of the method.

To summarize, structural seeds improved volume and sensitivity for detecting connectivity compared to traditional multivoxel seeds. For structural seeds, larger voxels (4mm vs. 3mm) improved connectivity except in posterior hippocampus; a larger smoothing kernel (10mm vs. 6mm) improved connectivity for both 3mm and 4mm voxels.

#### Effects of smoothing connectivity during repetitive tapping

For repetitive tapping, connectivity was examined using global analysis from both the left and right hippocampus, which improves sensitivity for detecting connectivity (36). We further examined smoothing kernel effects on connectivity from structural seeds during repetitive tapping, comparing effects of 6mm and 10mm smoothing kernels on 4mm isovoxels.

Structural seeds showed similar maxima locations in pre- and postcentral gyri with both smoothing kernels, but the extent of connectivity was invariably greater with the larger kernel (Table 7). Depending on the size of the smoothing kernel, a topographic arrangement of connectivity was discernible (Fig 4). In the central third of the hippocampus, a topography was observed along the medial / lateral axis with the 6mm smoothing kernel (Fig 4A left); connectivity from more lateral seeds (cyan and navy blue) extended progressively further posterior and superior. This topography was largely obscured with 10mm smoothing (Fig 4A, right), because larger clusters of connectivity from the middle seed (cyan) produced extensive overlap.

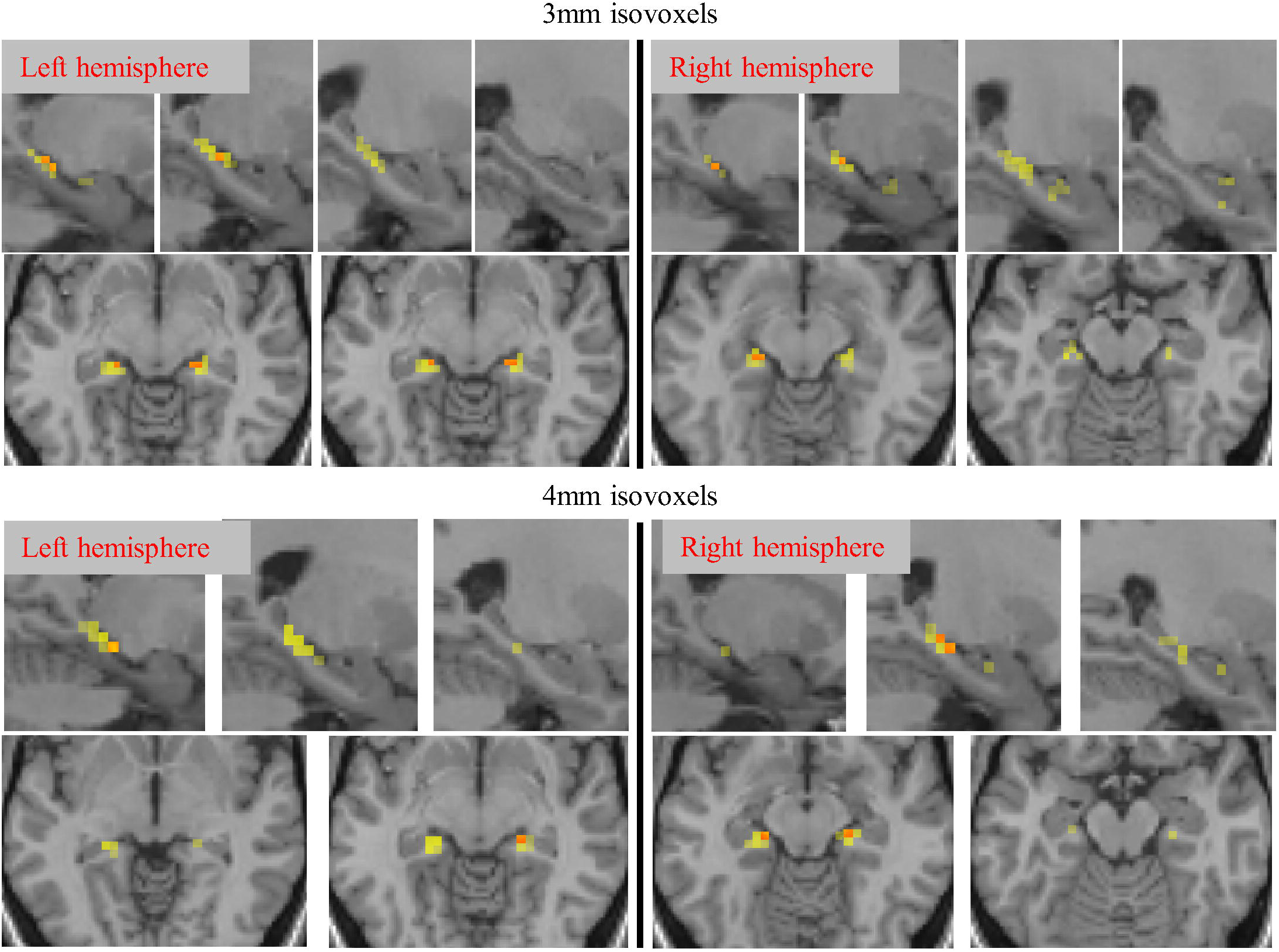

Along the anterior / posterior axis of the hippocampus, however, the topographic organization was more apparent with the 10mm smoothing kernel (Fig 4B). Connectivity for each seed was centered in the precentral gyrus, barely extending across the central sulcus into the postcentral gyrus. From the anterior hippocampal seed A1 (red), intense connectivity projected further anterolateral; the middle seed B1 projected superomedial (yellow), whereas the posterior seed C1 projected further medial and posterior (green). Because its clusters were smaller and spotty, this topography was indiscernible with the 6mm smoothing kernel.

Topography was also demonstrable from individual voxels with 10mm smoothing along both the medial-lateral and longitudinal axes (Fig 5). In the medial-lateral plane, the organization described above is seen in the pattern of overlapping connectivity maps (Fig 5A, left), as well as differential connectivity maps that illustrate which seeds generated the greatest amplitude of connectivity (Fig 5A, right). Along the longitudinal axis, less overlap was observed from individual voxels (Fig 5B), perhaps because of the greater distance between sampled voxels along this axis. From voxels in both planes, connectivity included both pre- and postcentral gyrus (Table 6).

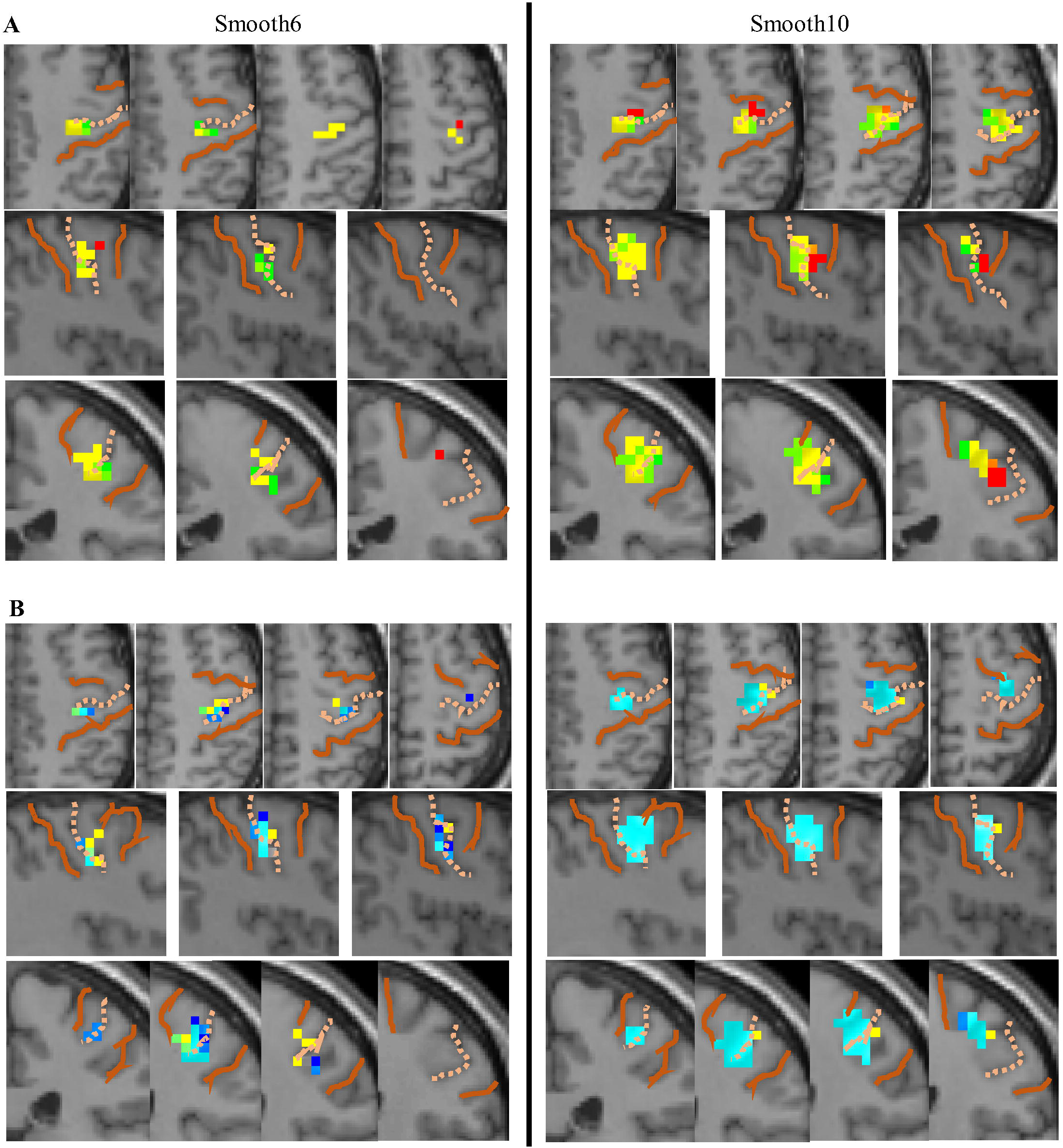

Precentral connectivity was more extensive in the right hemisphere during repetitive tapping (Fig 5), whereas postcentral connectivity was more extensive in the left hemisphere during sequential learning (Fig 4).

Overall, the 10mm smoothing kernel provided better sensitivity for structural seeds to demonstrate both hippocampal connectivity and topography.

### Hippocampal connectivity: Functional seeds

Functional seeds showed unilateral connectivity during group analysis. Tapseed^−^ generated right-hemispheric connectivity during the repetitive tapping task (Fig 6A, red), whereas memseed^−^ generated connectivity in the left hemisphere (Fig 6A, yellow). Smoothing kernels of 6mm and 10mm showed no differences in connectivity during repetitive tapping or in postcentral gyrus during sequence, whereas the extent of precentral connectivity during sequence learning was larger with the 10mm kernel for both memseed^−^ (2880 vs. 1600 mm^3^) and memseed^+^ (1728 vs. 320 mm^3^).

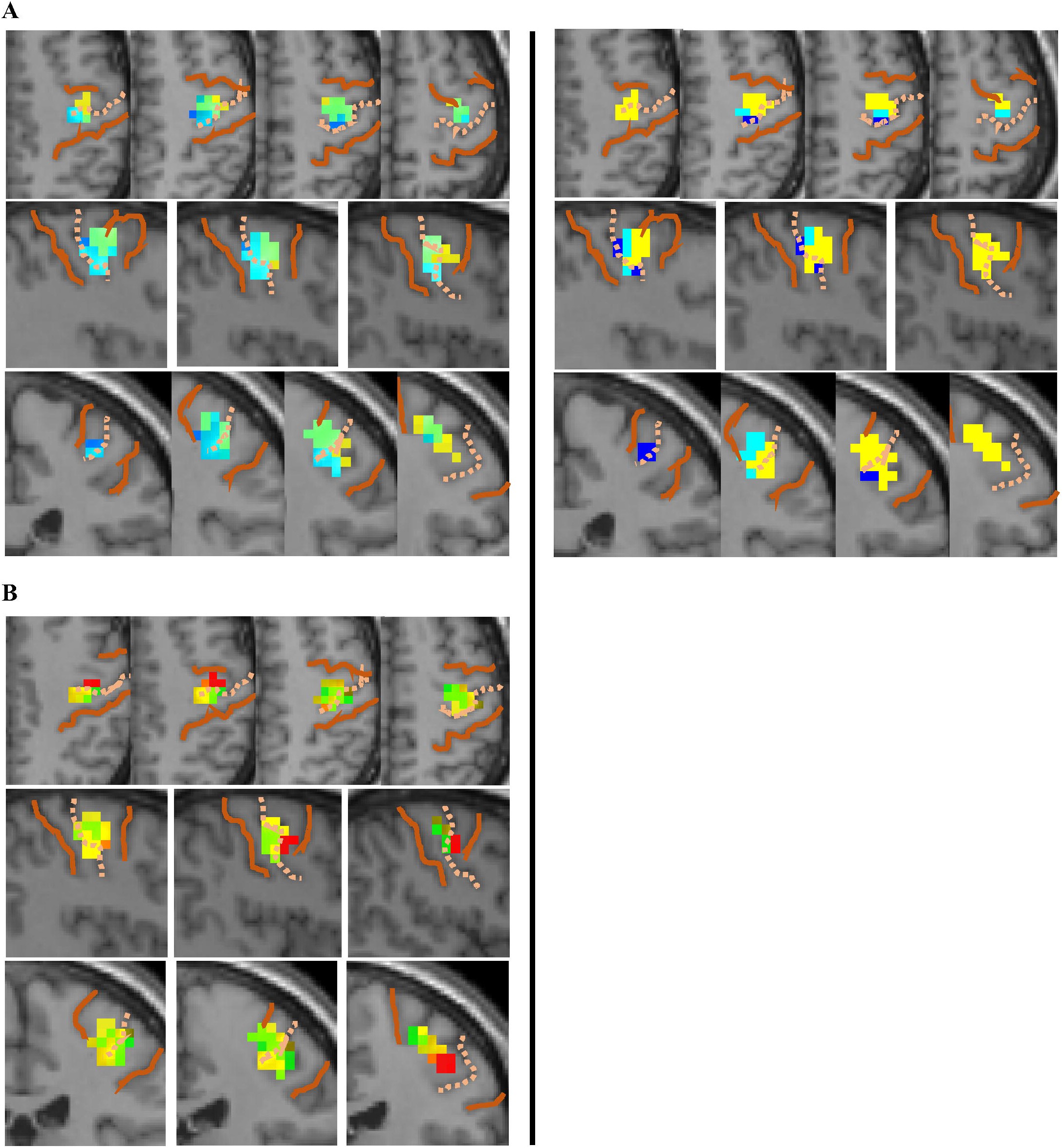

The 10mm smoothing kernel increased the intensity of connectivity as well as its extent. As noted earlier, a few hippocampal voxels individually showed greater motor activation with a 10mm vs. 6mm smoothing kernel; these voxels also showed significantly greater connectivity with 10mm smoothing (orange voxels, arrows) interspersed within the area of connectivity shown by functional seeds in both motor tasks (Fig 6B).

## Discussion

This study evaluated the effects of seed selection (structural vs. multivoxel vs. functional), voxel size (3mm vs. 4mm isovoxels), and smoothing kernels (6-10mm) on hippocampal connectivity with sensorimotor cortex; similarities to the effects of voxel size and smoothing kernels on SMC motor activation were also evaluated. Larger voxels modestly improved sensitivity for detecting activation in SMC; larger voxels also improved detection of connectivity from anterior regions of the hippocampus, although showed lesser sensitivity from the posterior third of the hippocampus. Larger smoothing kernels improved sensitivity for detecting both hippocampal activation and connectivity; this sensitivity allowed the topography of hippocampal connectivity to be mapped from single voxels as well as structural seeds. Although structural seeds showed greater sensitivity for connectivity than multivoxel seeds, they were more affected by the size of the smoothing kernel than functional seeds, which addressed individual variability by selecting voxels with maximal SMC connectivity within a hippocampal region.

### Effects of voxel size and smoothing kernel on cortical activation

Voxel size had modest effects on activation in this study. In SMC, larger voxels (4mm) showed a smaller statistical threshold and a larger activation volume during both repetitive tapping and sequence learning. These effects were small, detected through direct within-subject comparisons, whereas group comparisons of the mean showed no effect. Random-effects activation revealed only minor differences from voxel size in the area of activation along cluster borders; furthermore, no significant differences in contrast magnitude were observed, regardless of smoothing kernel. Although the larger 4mm isovoxels provide slightly better sensitivity for detecting cortical activation, this effect was modest.

The effect of smoothing kernel on cortical activation depended on the ROI. With the SMC mask, increases in the smoothing kernel from 6mm to 10mm resulted in the detection of much greater activation volumes; within the hand representation, this increase in activation volume was much less. This suggests the larger smoothing kernel more effectively reduced random variability (noise) from inactive regions within the SMC mask, reducing the mean difference in signal required to detect activation. This interpretation is consistent with observed decreases in the maxima t-value, contrast magnitude, and threshold.

During both individual and group analysis, the largest (10mm) smoothing kernel applied to 4mm isovoxels was most sensitive for detecting activation. This was not simply an overestimation of activation size from a large smoothing kernel, as suggested by others (15); global analysis across both tasks demonstrated an even larger, bilateral area of activation consistent with task behavior (36). With these processing parameters, the contrast estimate at the maxima from group analysis approached the mean value from individual analyses, suggesting this combination of voxel size and smoothing kernel provided the most accurate group representation of sensorimotor activation.

### Effect of smoothing kernel on hippocampal activation

Using conventional group analysis, the hippocampus did not show motor activation during repetitive movements or sequence learning. By using paired t-tests to compare motor activation across different smoothing kernels, greater activation was demonstrated with the 10mm smoothing kernel (compared with 6mm), evident with both 3mm and 4mm isovoxels. This differential activation occurred in the medial hippocampus, mostly within the central third of the hippocampus but extending slightly into the posterior third, most intense at the boundary between the B1 (central-medial) and C1 (posterior-medial) structural seeds.

During smoothing, local variability is minimized by recalculating the BOLD signal based upon peaks and their surrounding signal, fit to a Gaussian curve. When activity surrounding a 4mm isovoxel showing peak activity is symmetrical, both a 6mm and 10mm smoothing kernel utilize the magnitude of signal from a single voxel on each side in its calculation, differing only in the weighting provided from the adjacent voxel (3mm on each side of the peak 4mm-voxel using a 10mm kernel, and 1mm on each side of the peak 4mm-voxel with a 6mm kernel). Activity surrounding a peak voxel near the edge of gray matter is asymmetrical; the calculated peak coordinate is shifted toward the higher activity within gray matter, away from the edge (42, 43). Thus, differential activation from the larger smoothing kernel did not result from extending the smoothed signal outside hippocampal boundaries; instead, the 10mm kernel provided a better estimate of local activation by more fully incorporating information from nearby activity.

The region showing differential activation also showed more intense and widespread SMC connectivity following 10mm smoothing. Improved sensitivity for detecting local hippocampal activity with the larger smoothing kernel is thus useful for detecting its connectivity with other regions.

### Effect of voxel size and analytical approach on hippocampal connectivity

Because direct point-to-point comparisons is not possible, effects of voxel size were identified by comparing the volume of significant connectivity and maximal parameter estimates, using different methods and smoothing kernels. Using a 3×3 matrix, all hippocampal regions were sampled, thereby allowing detection of regional effects.

Traditionally, either the entire hippocampus or a multivoxel subregion is specified as a seed for connectivity analysis (44–47). Multivoxel seeds average hippocampal activity before looking for correlations elsewhere in the brain; this differs from our structural seed alternative, which identified the magnitude of correlations from each voxel within a hippocampal region, then averaged these connectivity maps. The method of analysis showed inconsistent effects on the maximal parameter estimate from a region; however, greater connectivity volumes were consistently observed from structural seeds in all regions, indicating this approach improves sensitivity for detecting connectivity. Indeed, multivoxel seeds failed to demonstrate significant connectivity in anterior hippocampus. Furthermore, the topographic organization of hippocampal connectivity only emerged with structural seed analysis.

In posterior hippocampus, 3mm voxels generated greater connectivity volumes than 4mm voxels, particularly in posteromedial hippocampus. This likely reflects both anatomical and functional properties of the hippocampus. The posterior hippocampus is thinner (four 4mm-voxels wide, compared with six in middle and anterior regions); furthermore, posteromedial hippocampus may be functionally more uniform. More lateral regions include positive connectivity, particularly in middle-to-anterior regions (e.g., see Fig S2 from Burman, 2019 (36)). A sharp transition between functionally distinct regions may thus be requisite for smaller voxels to demonstrate greater connectivity. By contrast, 4mm isovoxels generated greater connectivity volumes in anterior and central regions of the hippocampus, as well as generating larger parameter estimates at structural seed maxima (Fig 3 and Table 6).

In summary, structural seeds were more effective for demonstrating hippocampal connectivity. Except in posterior areas, perhaps due to a transition to a functionally distinct area, 4mm generated more extensive connectivity maps than 3mm voxels. In this study, a topographical organization to hippocampal connectivity was only evident with the structural seed analysis using 4mm voxels.

### Effect of smoothing kernel on hippocampal connectivity

This study showed different patterns of inverse hippocampal connectivity with SMC during sequence learning and repetitive tapping: the left hippocampus generated both pre- and post-central connectivity maxima in left SMC during sequence learning, whereas bilateral hippocampal seeds generated a single connectivity maximum in the right pericentral gyrus during repetitive tapping. The size of the smoothing kernel showed similar effects on connectivity during both tasks.

Structural seeds demonstrated larger clusters of SMC connectivity with the larger kernel in both motor tasks. A connectivity map from a structural seed represents the mean connectivity from every voxel within the designated region of hippocampus, so larger connectivity clusters reflect the effectiveness of the smoothing kernel in reflecting local regional activity (see also (48)).

With the 6mm smoothing kernel, a topographic pattern from structural seeds was evident along the medial-lateral axis of the hippocampus, but not the longitudinal axis, where small and disparate clusters were generated with no apparent topography. With the 10mm kernel, a topographic pattern was partially obscured along the medial-lateral axis, due to extensive overlap in connectivity from adjacent seeds, yet a topography along this axis was still evident from connectivity of individual voxels. Furthermore, the 10mm smoothing kernel was needed to demonstrate topography along the longitudinal axis of the hippocampus.

The size of the smoothing kernel may be a factor in detecting functional heterogeneity within the hippocampus. A resting state connectivity study using a small (3mm) smoothing kernel failed to find functional heterogeneity along the anterior-posterior axis (49), whereas a study of connectivity from the hippocampus to cortical regions during episodic memory found functional heterogeneity along the same axis using a larger (5mm) smoothing kernel (25).

In summary, the 10mm smoothing kernel was advantageous for demonstrating hippocampal topography in its connectivity with SMC.

### Topography of hippocampal connectivity

Similar to previous studies, hippocampal connectivity in this study showed a topographical organization along both the long anterior-posterior axis (50–53) and medial-lateral axis (50).

In the current study, hippocampal connectivity with SMC was preferentially seen within the SMC hand representation for both motor tasks, with location and laterality similar to group activation by the tasks. Group activation and inverse connectivity were both limited to the left SMC during sequence learning (reflecting right-handed movements) and the right SMC during repetitive tapping (despite bilateral hand movements). Furthermore, activation and connectivity both covered the breadth of the postcentral gyrus within the hand representation during sequence learning, whereas activation and connectivity were both centered in precentral gyrus during repetitive tapping, with postcentral connectivity limited to the region adjacent to the central sulcus. Whether these task differences arose from differences in behavioral requirements, hand dominance (right-handed subjects), or hemispheric differences in motor function is unknown.

The role of hippocampal topography to SMC function is unclear. Within the hand area of the precentral gyrus, overlapping but distinguishable regions are involved in wrist and individual finger movements (32, 54–56); multiple representations for each digit may differentially reflect flexion and extension movements (35) or different degrees of movement complexity (57). Similarly, somatotopy for individual fingers in postcentral gyrus are overlapping but distinguishable, with multiple representations across the anterior-posterior width of the gyrus (58–60). Methods here did not allow mapping onto individual finger representations, or even flexion vs. extension, so the functional significance of the hippocampal topography observed here is unclear.

Different cognitive functions have been suggested for anterior vs. posterior hippocampus, differences such as encoding vs. recall (61, 62), internal vs. external attention (63), gist/conceptual vs. detailed/spatial information (64–66), and pattern completion vs. separation (67). In the current study, the topographical pattern of connectivity observed from anterior vs. posterior hippocampus is inconsistent with these roles, particularly as the repetitive tapping task required paced, volitional movements without learning or memory recall (36). Topographical connectivity with different regions within the pre- or postcentral hand representation suggest a role more directly involved in motor control, perhaps coordinating cognitive and motor functions to facilitate and suppress muscle actions required for the correct timing of finger movements (37, 56).

### Practical considerations for future studies

Many hippocampal studies have used small voxels and smoothing kernels, believing these processing parameters more accurately reflect true hippocampal activity. The current study suggests this may not always be the case, particularly when examining hippocampal connectivity with cortical areas. This study specifically examined the effects of voxel size and smoothing kernel on hippocampal activation and connectivity with SMC; however, results should apply to other hippocampal studies as well, as observed effects from changes in processing parameters are consistent with previous studies (as noted earlier).

Results also indicate greater sensitivity can be obtained with structural seeds, rather than conventional multivoxel seeds. For structural seed analysis, connectivity is calculated from each voxel of the hippocampus, then a mean connectivity map is calculated from all voxels in a region. This approach can map connectivity from the entire hippocampus, particularly advantageous when the precise location of a functional region is unknown.

Paired t-tests comparing cortical activation with 4mm vs. 3mm isovoxels showed lower thresholds and larger volumes of activation from larger voxels, with the additional volume expanding the edges of an activation cluster. Similarly, larger smoothing kernels decreased the threshold and increased the volume of cortical activation. Even small differences in activation between 8mm and 10mm smoothing kernels were significant during sequence learning, although these differences did not reach significance for 4mm voxels during repetitive tapping.

Parameters optimal for detecting SMC cortical activation also improved hippocampal connectivity (10mm smoothing kernels and 4mm voxels, except from the posterior third of the hippocampus). This similarity suggests better detection of cortical activity might allow better correlation with hippocampal activity. Larger smoothing kernels also improved the signal-to-noise ratio with local hippocampal activity, however, as shown in direct comparison between 10mm vs. 6mm smoothing kernels. Furthermore, localized areas with elevated hippocampal activity from the 10mm smoothing kernel also showed elevated connectivity with SMC, indicating this activation was functionally relevant. The effectiveness of these parameters thus reflect both cortical and local hippocampal properties.

Because the size and structure of the hippocampus is affected by a wide array of factors, including age, exercise, depression, and stress (68–73), its functional localization must be variable. Functional seeds sought to reduce effects of individual variability during group analysis. Restricted to structural regions showing significant connectivity during group analysis, the voxel showing maximal connectivity within the entire SMC was identified. Using this ROI during seed selection, the selection of functional seeds did not bias the outcome; connectivity maxima used for seed selection were scattered, located in both hemispheres and often outside the hand representation (36). These functional seeds nonetheless showed selective connectivity within the hand region during group analysis. The current study showed that the sensitivity of functional seeds was minimally affected by the size of the smoothing kernel, suggesting smoothing mitigates the effects of individual variability during group analysis, but also that sensitivity for demonstrating connectivity with non-optimal parameters can be improved by identifying functional seeds.

## Conclusions

This study shows that processing fMRI data with a larger voxel size (4mm) and smoothing kernel (8-10mm) can improve sensitivity to hippocampal activity and its connectivity with cortical areas; optimizing these processing parameters uncovered topography in hippocampal connectivity along two axes. Structural seeds that represent the mean connectivity from all voxels in a hippocampal region provide better sensitivity than traditional multivoxel seeds, and have the additional advantage of mapping connectivity throughout the hippocampus.

## Acknowledgments

The author wishes to thank the Center for Advanced Imaging (CAI) at NorthShore University HealthSystem for its administrative support.

